# *In Vivo* Monitoring of Rare Circulating Tumor Cell and Cluster Dissemination in a Multiple Myeloma Xenograft Model

**DOI:** 10.1101/516641

**Authors:** Roshani Patil, Xuefei Tan, Peter Bartosik, Alexandre Detappe, Judith Runnels, Irene Ghobrial, Charles P. Lin, Mark Niedre

**Author notes:** RP and XT were equal contributors to this work.

## Abstract

We recently developed ‘Diffuse *in vivo* Flow Cytometry’ (DiFC), a new pre-clinical research tool for enumerating extremely rare fluorescently-labeled circulating cells directly *in vivo*. In this paper, we developed a green fluorescent protein (GFP) compatible version of DiFC, and used it to non-invasively monitor the circulating tumor cell (CTC) burden over time in a multiple myeloma disseminated xenograft model. We show that DiFC allowed counting of CTCs at estimated concentrations below 1 cell per mL in peripheral blood with a negligible false alarm rate. DiFC also revealed the presence of CTC clusters in circulation to our knowledge for the first time in this model, and allowed us to calculate their size, kinetics, and frequency of shedding. We anticipate that the unique capabilities of DiFC will have many applications in the study of hematogenous metastasis, and as a powerful complementary methodology to liquid biopsy assays.

## 1. Introduction

Circulating tumor cells (CTCs) are understood to be important in the development of hematogenous cancer metastasis. Their presence and numbers are known to be associated with prognosis for many cancer types (1,2). As such, CTCs are a major focus of clinical and preclinical research (3–5). CTCs are exceedingly rare – fewer than 1 CTC per mL of peripheral blood is associated with poor overall survival. Enumeration of CTCs is normally performed by drawing and analyzing peripheral blood samples, wherein target cells are isolated using methods including flow cytometry, size-based cell separation, immuno-magnetic separation, and microfluidic capture (6).

Although ‘liquid biopsy’ tools remain the workhorse of pre-clinical CTC research they remain far from optimal. In the context of longitudinal studies of rare CTC populations they have a number of known limitations (7,8): for survival experiments, blood collection is limited to about 10% of the peripheral blood volume every two weeks (9), which may result in serious inaccuracies in estimating the CTC burden due to sampling error (10,11). In some cases involving rare cells, mice must be euthanized so that the entire peripheral blood volume may be analyzed, eliminating the possibility of longitudinal study over time. Moreover, limited temporal sampling makes it difficult to observe the kinetics of CTC shedding, which may vary significantly over timescale of days and (as we show) even minutes. Moreover, blood samples are known to degrade rapidly after removal from the body (12), and the process of drawing blood can trigger a stress response in the animal (13).

In addition to single CTCs, CTC clusters (CTCCs; also called circulating tumor microemboli; CTM) are multicellular groupings of CTCs that exhibit a mixture of epithelial and mesenchymal properties, and may include stromal (14) and other non-tumor cells (15,16). CTCCs are even more rare than individual CTCs, but are known to have significantly better survivability in circulation and higher (~100X) metastatic potential. Despite their recognized importance, much is still not understood about CTCCs, including the mechanisms for their survival advantage *in vivo*, as well as their abundance, kinetics of shedding, and composition. It has been suggested that a key reason for this is that nearly all CTC isolation methods are not designed to detect clusters and therefore may result in their dissolution or loss (15). This may contribute to a general underestimation of their importance in the CTC research community, and has also driven development of microfluidic systems specifically designed to isolate clusters in recent years (17).

Our team recently developed a new small animal research tool termed ‘Diffuse *in vivo* Flow Cytometry’ (DiFC) (18). DiFC is a variant of ‘*in vivo* flow cytometry’ (IVFC) which is a general class of instruments for counting circulating cells in the bloodstream without drawing blood samples (7,8,19). The distinguishing feature of DiFC compared other IVFC methods is that it uses diffuse light to detect fluorescently-labeled circulating cells in large blood vessels such as the ventral caudal artery (VCA) in the tail of a mouse. DiFC uses a dual-optical probe configuraiton placed over a large vascular bundle. We previously showed that DiFC allows interrogation of hundreds of microliters of circulating blood per minute in nude mice (18). In contrast, microscopy-IVFC samples small blood vessels in the mouse ear where blood flow rates are on the order of 1 μL/min (8). In our previous work, DiFC used red and near infrared (NIR) fluorescent dyes, because of the minimal attenuation of NIR light in biological tissue (20). However, this limited the use of DiFC to cells expressing NIR fluorescent proteins (FPs) (21) which are significantly less common in biomedical research than visible FPs such as the green fluorescent protein (GFP) (8,13,22).

The novelty of the present work is three-fold. First, we developed a blue-green version of DiFC suitable for use GFP, thereby greatly expanding the utility of DiFC. As we show, we discovered that while blue-green light exhibits somewhat less favorable tissue optics compared to NIR light (20), the relative loss of signal for DiFC was actually modest since that it operates in relatively superficial tissue volumes (< 2mm). The instrument was built on a compact optics cart for ease of movement between sites.

Second, we used our GFP-DiFC system to monitor the growth and vascular spread of multiple myeloma (MM) in a disseminated xenograft model (DXM) in mice (23). MM is a hematological malignancy that is believed to originate at a single site in the bone marrow niche, and then continuously disseminate by the circulatory system. Hence, MM is of significant interest to researchers, both as a disease and as a model of cancer metastasis (24). We show that the unique capabilities of DiFC allowed us to follow development of MM longitudinally in individual mice for up to 36 days after inoculation, at CTC burdens below 1 cell per mL with negligible low false alarm rate (FAR).

Third, DiFC revealed the presence of MM CTC clusters in circulation in this xenograft model to our knowledge for the first time. Although there is some prior evidence that MM grows in clusters in the bone marrow (23) and may be found in peripheral blood in patients (24), very little is known about MM CTC clusters in general. The capabilities of DiFC allowed us to quantify the frequency of CTCC shedding and estimate the size of clusters directly in vivo during disease development. These data are extremely difficult to obtain with liquid biopsy methods, and illustrate the value of complementary *in vivo* techniques such as DiFC in the study of hematogenous metastasis.

## 2. Methods and Materials

### 2.1 DiFC Instrument

The schematic of the DiFC instrument is shown in ***figure 1a***. The system is a similar design to the previously-reported NIR system (18), but uses optical components compatible with GFP. Specifically, the light source was 488 nm DPSS laser (DL488-150; Crystalaser LLC, Nevada), the output of which was filtered with a cleanup band-pass (BP-x) with 488/10 nm (ZET488/10x; Chroma). These were coupled into the ‘source’ fiber of the fiber probes using lens-fiber couplers (FC-x) with 532nm anti-reflection coating (F240SMA-532; Thorlabs). Two filters were mounted directly to the tip of the fiber bundles: a central band-pass filter (BP-f) at 488/5 nm to filter the excitation light, and an outer detection ring shape long-pass filter (LP-f) at 503 nm to collect the fluorescence light. The output of the collection fiber bundles (8 per probe) were terminated on a second set of lens fiber couplers (FC-m), and then filtered with interference filters (BP-m) at 535/50 nm (ET535/50m; Chroma). The band-pass filters were identical here, but could be different in the future, for example to allow measurement of 2-fluorophores. Four current-output photomultiplier tubes (PMTs; H6780-20, Hamamatsu, New Jersey) were powered by a voltage supplies (C10709, Hamamatsu). The output of each PMT was amplified with low-noise current pre-amplifiers (PA; SR570, Stanford Research Systems, Sunnyvale, CA) with 300 Hz low pass filter, and then digitized with a multi-function data acquisition board (USB-6212 BNC; National Instruments, Austin, TX). The entire setup was mounted was mounted on an optics cart (POC001, Thorlabs), so that it could be easily between sites (***fig. 1b***).

**Figure 1.**
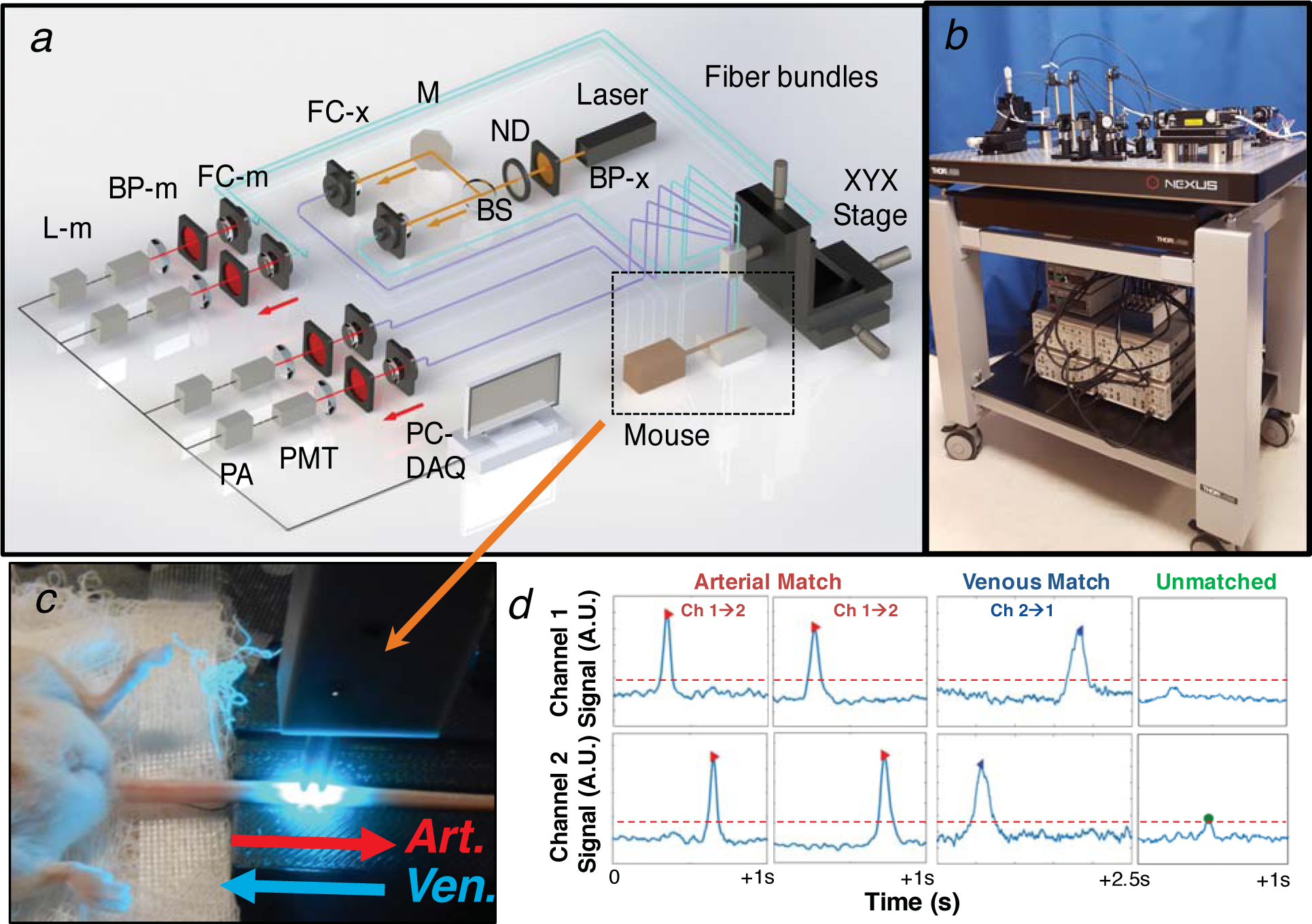
(a) DiFC instrument schematic (see text for details). (b) The DiFC system was mounted on an optics cart that could be moved easily between sites. (c) The fiber bundles were placed on the ventral surface of the mouse tail, approximately above the ventral caudal bundle. (d) DiFC allows detection and discrimination of circulating MM GFP-labeled cells moving the arterial and venous directions. Peaks that are not measured on both channels are presumed to be moving in smaller blood vessels or are due to noise, which are subsequently discarded from the analysis.

### 2.2 DiFC Data Acquisition

Mice were held under inhaled isofluorane during DiFC scanning to prevent movement, and were kept warm using two heating pads placed under the body and over the exposed area of the tail. The DiFC fiber probes were separated by 3 mm and were placed approximately over the large VCA on the ventral side of the tail (***fig. 1c***). Mice were scanned with DiFC twice per week for 45 minutes at 2000 samples per second. The first 10 minutes of acquisition data were discarded since we often observed transient effects which we attributed to the warming of the tail with the heating pad (i.e. a transient increase in blood flow and CTC count rate).

### 2.3 DiFC Data Analysis

DiFC data was analyzed as we described previously (18) with a few minor differences. Data was pre-processed by summing the data from the two PMTs per bundle, applying background subtraction, and then applying a 5 ms moving average filter. Data from the two channels were then analyzed using a two-step procedure. In the first step, ‘cell candidates’ were detected in the data using a set threshold of 75 nA (this was empirically selected to yield a low FAR). In the second step, candidates were matched in either the forward or reverse directions, according to similarities in the peak width, amplitude, and separation in relation to the estimated cell speed. This allowed us to distinguish cells moving the forward (arterial) and reverse (venous) directions, and to distinguish ‘unmatched’ peaks as false alarm signals due to electronic noise or motion artifacts (***fig. 1d***).

A major new result in this manuscript versus our previous work is the detection of CTC cluster-like signals with DiFC. These CTCC-like signals were characterized as having large amplitude and width relative to individual cells, indicating the presence of a grouping. We automated detection of these as having minimum width of 200 ms and minimum amplitude of 4 times the detection threshold (300 nA). We also estimated the equivalent number of cells in a cluster by dividing the amplitude of the cluster by the mean of a single cell.

### 2.4 Multiple Myeloma Disseminated Xenograft Model

All mice were handled in accordance with Northeastern University’s Institutional Animal Care and Use Committee (IACUC) policies on animal care. Animal experiments were carried out under Northeastern University IACUC protocol #15-0728R. All experiments and methods were performed with approval from, and in accordance with relevant guidelines and regulations of Northeastern University IACUC.

We used MM.1S multiple myeloma cells that were genetically modified to carry GFP, firefly luciferase, and neomycin genes (MM.1S.GFP.Luc cells). These cells were originally described by Dr. Rosen at Northwestern University. Cells were authenticated by an external service (Bio-Synthesis Inc., Lewisville, TX) to verify their MM.1S lineage.

We used 8-week-old male SCID/Bg mice (Charles River) tail vein injected (*i.v.)* with 5 x 10^6^ MM.1S.GFP.Luc cells suspended in 200 µL PBS (N = 8) or PBS injected controls (N = 4). SCID mice required removal of the hair on the tail region with depilatory cream (Nair). We also used clear imaging gel (Ultrasound and Laser Gel #4963, McKesson Medical-Surgical Inc. Richmond, VA) applied to the skin surface to facilitate optical coupling. Control mice were always scanned on the same day as MM bearing mice, to account for the possibility that the background noise properties of the instrument may change day-to-day.

Mice were grown in two separate cohorts of N = 4 tumor bearing and N = 2 control each. For reasons described in detail below, for the first cohort (‘cohort 1’; C1), mice were followed up to 31 days after injection. For the second cohort (‘cohort 2’; C2), mice were followed up to 36 days after injection.

### 2.5 Bioluminescence Imaging

As a secondary method for tracking MM growth, bioluminescence imaging (BLI) was performed weekly with a commercial IVIS Lumina II imaging system (Caliper Life Sciences, now Perkin Elmer, Waltham, MA). Mice were injected i.p. with 150 mg/kg of D-luciferin (Perkin Elmer) 10 minutes prior to imaging. The image exposure time was one minute.

### 2.6 Flow Cytometry

For mice in C1, we drew blood samples for analysis with flow cytometry on day 24 (200 µL) and day 31 (0.5-1 mL, terminal) to verify the presence of MM CTCs in the blood. We counted GFP+ cells in drawn blood samples using the green channel of a commercial flow cytometer (Attune NxT, Thermo Fisher). The gating and counting process was as follows: We first performed FC on cultured MM.GFP+ cells in suspension to determine the appropriate SSC-FSC gates (***Supplemental Figure S1a***) and distribution of 530/30 nm blue fluorescence (***Fig. S1b)*** for the cells. We used the mode fluorescence intensity of Dragon Green 2 microspheres (DG2; Bangs Laboratories Inc., Fisher, IN) as a reference and a fluorescence threshold for cell counting as shown by the dotted red line. Blood samples were pre-processed by first lysing the RBCs with a lysis buffer (420301, Biolegend, San Diego, CA) according to the manufacturer’s instructions. An example SSC-FSC plot for a blood sample (taken on day 31) is shown (***Fig. S1c)*** as well as the blue fluorescence histogram (***Fig. S1d)***, wherein most signal was non-fluorescent debris. Application of the threshold (which was defined as the intensity of DG2 microspheres for all samples) allowed us to count MM.GFP+ cells. These numbers were divided by the volume of drawn blood to estimate the concentration of MM cells *in vivo*.

### 2.7 Blood Smear Preparation and Imaging

For mice in C2, we terminated the experiments on day 36 after injection. The aim was to verify the presence of CTC clusters in vivo, which as we show was suggested by our DiFC data. We drew and collected 750 µL of blood in EDTA tubes and euthanized the mice. We created 30 blood smears per mouse by pipetting 6 µL of blood on a glass microscope slide (Fisherbrand Colorfrost microscope slides with clipped corners, Fisher). The cell monolayer area of each slide was imaged with an upright Carl Zeiss microscope with an HXP 120C light source using the bright field channel and the eGFP channel to detect the presence of MM.GFP+ CTCs and CTCCs. Examples images were cropped to 65 x 65 µm^2^ to show example cells and clusters of interest. Overlays were performed using the ‘merge channels’ function in ImageJ.

### 2.8 Monte Carlo Simulations

We used an open source, GPU-accelerated Monte Carlo program (Monte Carlo eXtreme) to compute the detection sensitivity functions for DiFC (25). We modeled the tail as a homogenous 4 mm diameter, 4 cm long cylinder, with voxel size of 250 µm^3^. We used literature values (20) for optical properties including scattering coefficient (*µ*_*s*_), absorption coefficient (*µ*_*a*_), at the excitation (*ex*) and emission (*em*) wavelengths, and the anisotropy coefficient (g). These were as follows. For NIR wavelengths: *µ*_*s-ex*_ = 22 mm^-1^, *µ*_*s-em*_ = 20 mm^-1^ *µ*_*a-ex*_ = 0.002 mm^-1^, µ_a-em_ = 0.0015 mm^-1^, *g* = 0.9. For blue-green (GFP) wavelengths: *µ*_*s-ex*_ = 40 mm^-1^, *µ*_*s-em*_ = 38 mm^-1^, *µ*_*a-ex*_ = 0.02 mm^-1^, *µ*_*a-em*_ = 0.02 mm^-1^, *g* = 0.9.

## 3. Results

### 3.1 MM Disseminated Xenograft Model

We used an MM disseminated xenograft model (DXM) as has been described by our team previously (23). MM.1S.Luc.GFP cells were injected *i.v.* in SCID/Bg mice. MM cells rapidly home to the bone marrow, and then steadily proliferate throughout the skeleton over time. MM cells are eventually observed in circulation in peripheral blood, mimicking the clinical course of the disease. As shown in ***figures 2a-c***, we verified the growth of MM in our mice by weekly bioluminescence imaging (BLI). As expected, MM grew in a diffuse pattern in the skeleton, primarily in skull, spine and hips. Small amounts of BLI above background were observed as early as day 23, with significant disseminated growth by day 30. Significant variability in MM growth was observed between mice (***fig. 2d***), which may be due to differences in the efficiency of the tail vein injection of MM cells. In one mouse (M3C2) no development of MM was observed by BLI (or by DiFC; see below), however, we chose to keep it in the analysis since it served as an additional blind control.

**Figure 2.**
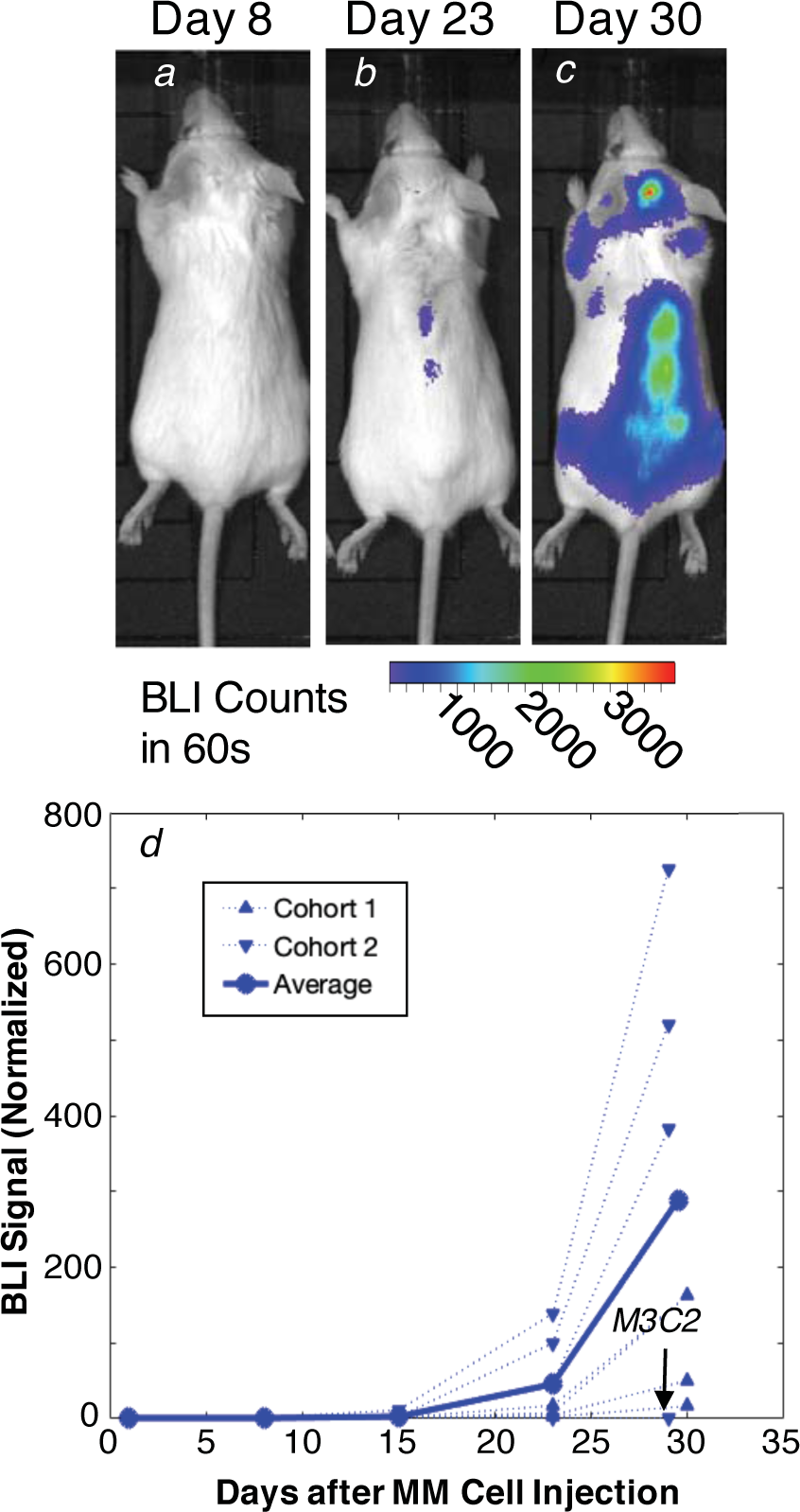
We performed BLI imaging weekly for all mice for 1 month following tail vein injection of MM.1S.GFP.Luc cells. (a-c) Luminescence increased as the MM disease grew in the bone marrow, which was first observable in small areas of the (b) spine by day 23. (c) By day 30, diffuse patterns of MM growth were observable in the skull, spine and hips. (d) This general pattern was observed for all inoculated mice except for one (M3C2), with significant inter-experimental variability between mice.

### 3.2 DiFC Performance at Blue-Green Wavelengths

We built a blue-green DiFC system for use with GFP expressing cells and transgenic mice models (13,26). Although we anticipated that DiFC might perform poorly at blue-green wavelengths where light attenuation is relatively high (20), Monte Carlo simulations indicated that the expected attenuation with GFP was only 20% higher than at NIR wavelengths in the 1-2 mm DiFC detection depth. As such, although DiFC relies on highly scattered light, optical attenuation was not as serious a problem as with whole animal imaging (27). Subsequent analysis showed similar modest expected losses with wavelengths corresponding to mCherry and Yellow Fluorescent Protein (YFP), so that we anticipate that DiFC systems corresponding to these wavelengths could be developed in the future.

Additional noise due from tissue autofluorescence at blue-green wavelengths was another potential concern. However, we tested the blue-green DiFC design in SCID mice and found that the background was about 10 μA with 20 mW of power at the sample, which was similar to our NIR system. Likewise, the noise after background subtraction was less than 25 nA, which was actually slightly lower than our NIR system. We scanned 4 control mice, twice weekly with DiFC after the initial sham (PBS) injection, giving 17.5 cumulative hours of DiFC scanning on non-tumor bearing mice. We did this as a test of the stability if the DiFC system, i.e. since no GFP+ cells were present, any ‘detections’ were false alarms. On average, we found that the false alarm rate (FAR) was extremely low over all DiFC sessions, with an overall average FAR of 0.016 per minute (or one every 70 minutes). An example 10-minute trace from a control mouse is shown in ***figure 3a***.

**Figure 3.**
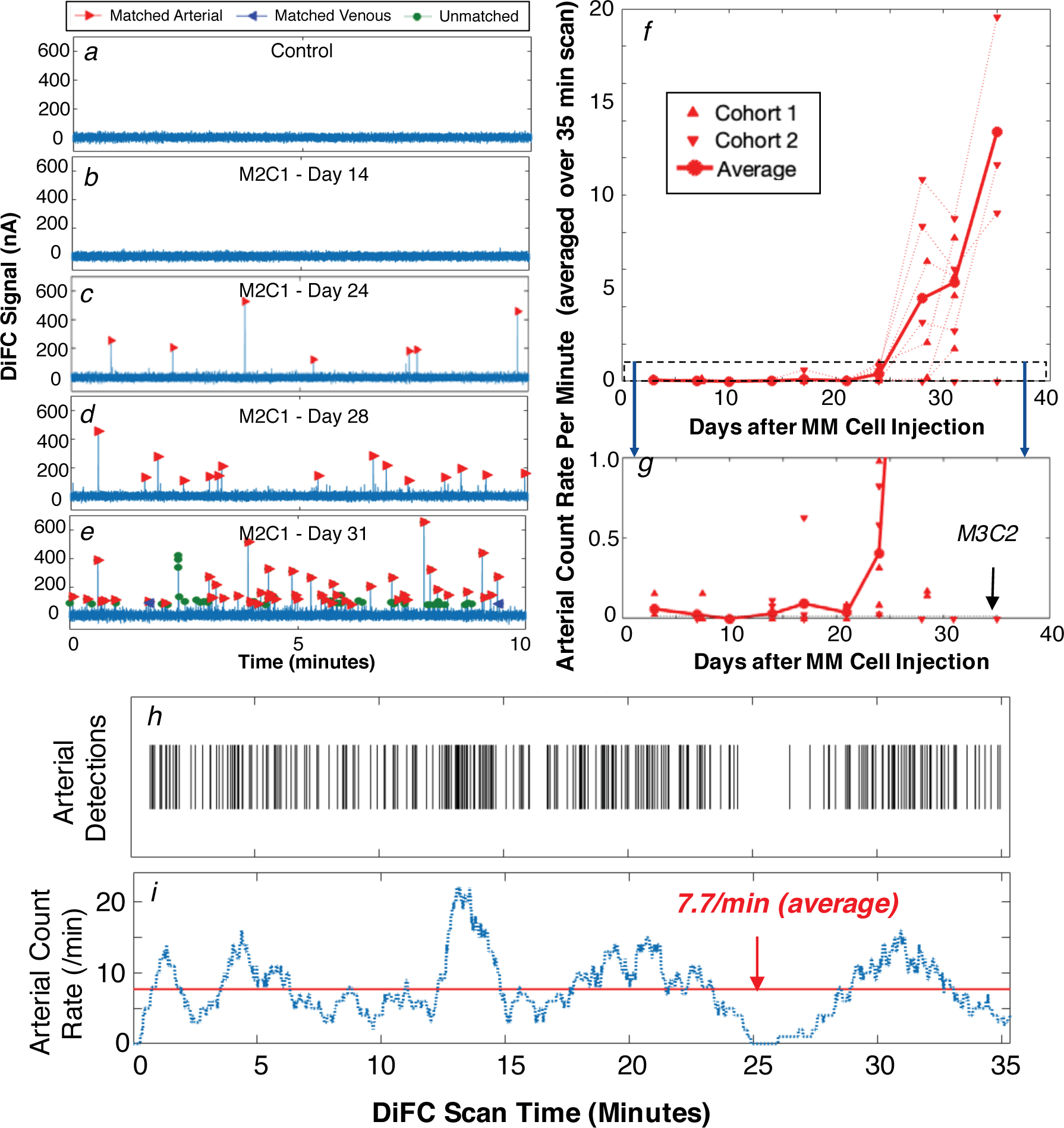
We performed DiFC scanning twice per week during development of the xenograft model. (a) An example 10-minute DiFC sequence from a control (PBS injected) mouse is shown, illustrative of the low false alarm rate of the DiFC system. (b-e) As the MM disease progressed, GFP+ MM cells were observable in circulation with increasing frequency. (f) The mean count rate in the arterial direction for all mice is shown, showing growth over the course of the disease, as well as significant inter-experimental variability. (g) An expanded view of (f) for mean DiFC-count rates between 0 and 1 counts/min. DiFC allowed detection of very low numbers of circulating cells above the FAR (dotted line, 0.016/min). As with BLI, one mouse (M3C2) failed to show signs of MM growth. (h) Raster plot showing the detection times of arterial-matched CTCs for mouse M2C1 on Day 31 during DiFC scanning, where each vertical line represents one detected cell. (i) The average count rate for this scan was 7.7 per minute (red line), but significant variability was observed when considering a 60-second moving average window (blue dotted line).

### 3.3. CTC Dissemination During MM Growth

We scanned the MM-bearing mice with DiFC twice per week after injection. Example 10-minute sequences of DiFC data from a single mouse (mouse 2, cohort 1; M2C1) from day 14 to day 31 is shown in ***figs. 3b-e***. As we showed previously (18), DiFC allows us to distinguish cells moving in the arterial (red markers) and venous (blue markers) directions.

We considered the average (mean) CTC count rate detected in the arterial (VCA) direction during the progression of the disease, as summarized in ***fig. 3f*** for all 8 MM-inoculated mice we studied. As shown in ***fig. 3g*** small numbers of arterial matched GFP+ CTCs were observed during the first few weeks (<1 count/minute). By day 21, rapid growth in MM CTC numbers was observed in all but one inoculated mouse. This was the same mouse (M3C2) that failed to show development of MM on BLI, again, most likely due to a failed intravenous injection of cells. As with BLI, significant inter-experimental variability between the mice was observed.

Another interesting feature of the data is that significant *short-term* variability in the DiFC count rate was observed, with periods of significantly lower or higher detection rates. For example, ***Fig. 3h*** shows the full 35-minute timeline of CTC detections for M2C1 on day 31, where each vertical line represents the detection of an arterial-matched cell. ***Fig. 3i*** shows the mean count rate for the scan (red line; 7.7 per minute) and the moving average over 60-second intervals (blue dotted line), where the count rate varied from 0 to 22 per minute during scanning. This significant variability was always observed for the mice in this study and may be expected in part from statistical effects, particularly with small time samples and rare cells (10,11). The implications of this are discussed in more detail below.

DiFC also allowed us to measure the speed of cells in the arterial direction by analyzing the arrival time between the two fiber probes, which were separated by 3 mm (***fig. 1d***). For these mice, the average speed of cells in the arterial direction was 26.5 mm/s. We note that this was significantly slower than we measured in nude mice previously (18), where the average arterial cell speed was 112 mm/s. We attribute the difference to the physically smaller size of the tails (and presumably VCA) of SCID/bg strain compared to nude mice. The measured peak *widths* for cells in the arterial direction was 39 ms in SCID mice, which is consistent with the slower speed, i.e. slower moving cells take longer to move through the ~1 mm DiFC field-of-view.

### 3.4 MM CTC Clusters

Interestingly, DiFC scanning also revealed the presence of large, irregular-shaped signal features such as those shown in ***Figures 4a-d.*** These were significantly wider (temporally) and higher (amplitude) than individual matched cells. We hypothesized that these signals originated from MM CTC clusters in the blood. As shown in ***fig. 4e***, CTCC signals were observed soon after single CTCs were detectable (day 24), and appeared with increasing frequency as the disease progressed to a maximum of about 2 per minute. We also note that these CTCC-like signals were never observed in control mice, nor were they observed in our previous experiments involving mice intravenously injected with cultured MM cells (18,28), implying that these originated from groupings of MM cells growing in the bone marrow niche that shed in to circulation.

**Figure 4.**
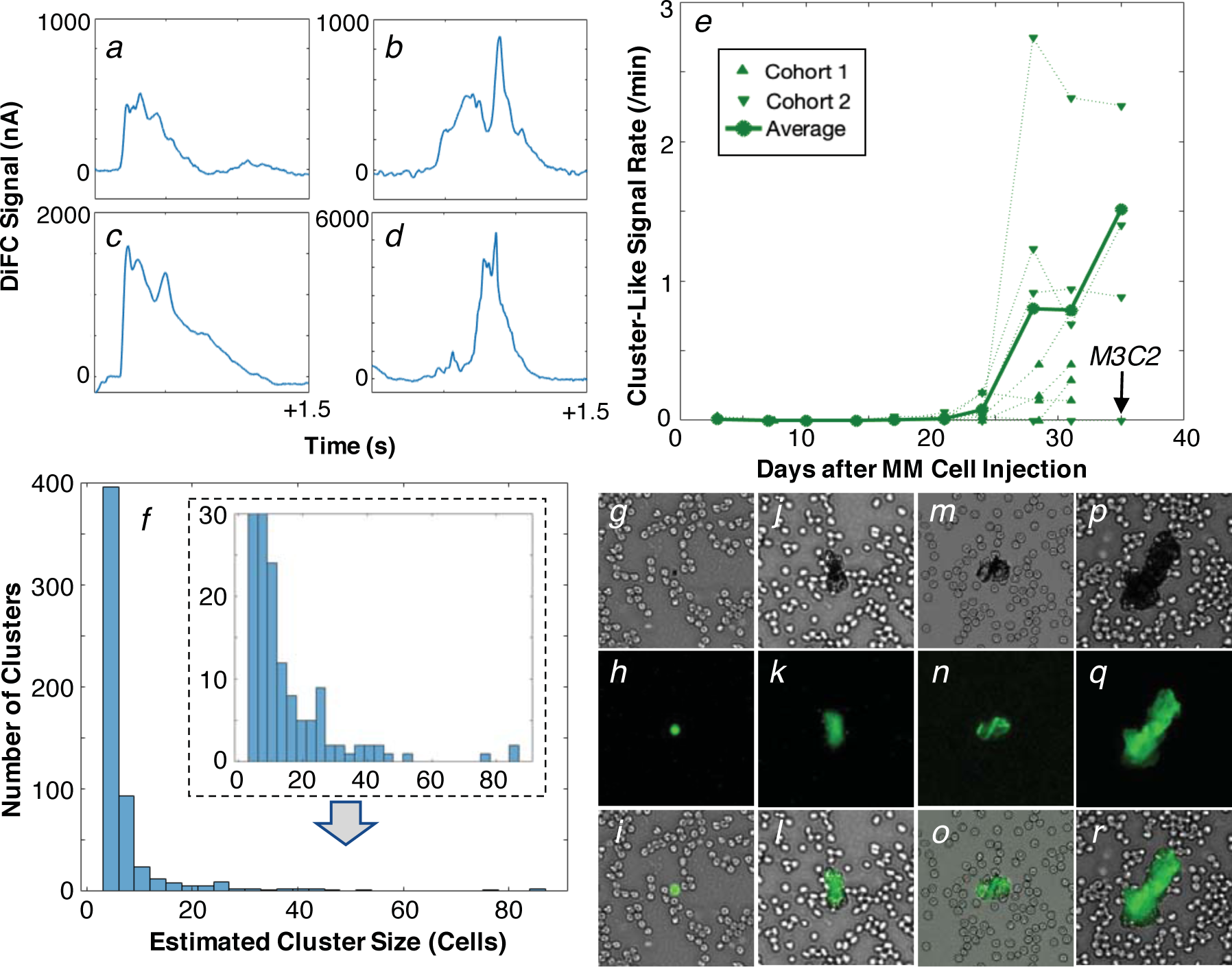
(a-d) DiFC also showed large, irregularly shaped signals (relative to individual cells), which we hypothesized were due to the presence of CTC clusters. (e) These were observed with MM-injected mice only (never controls), and appeared approximately as soon as single CTCs were observed in circulation at a rate of approximately 10% of the arterial count rate. (f) Analysis of the peak amplitudes allowed us to estimate the sizes of the clusters, which were frequently fewer than 10 cells, but individual (inset) clusters were very large, with dozens of cells. (g-r) Example white light, GFP fluorescence, and merged (overlay) blood smear micrographs from MM.1S bearing mice sacrificed on day 36 after injection. Most observed MM GFP+ cells were single cells (g-i), but small numbers of CTC clusters (j-r) were also observed as predicted from our DiFC measurements. Each image is 65 x 65 µm^2^.

We analyzed the pulse amplitude of the CTCC-like signals relative to single peaks, and estimated the equivalent number of cells in clusters as shown in ***Fig. 4f***. Using this, most MM CTCCs were usually estimated to be smaller than 10 cells, with mean and median estimated sizes of 6.4 and 4 cells, respectively. However, a small number large CTCC-like peaks were also observed with estimated sizes approaching 100 cells (***fig. 4f inset***). We did not use the width of the peak in estimation of the CTCC sizes, since, unlike microscopy-IVFC (29), even large clusters of cells are much smaller than the ~1 mm DiFC field of view. Therefore, we attribute the broad temporal width of CTCCs relative to individual CTCs to significantly slower speed of motion in the blood vessel, as opposed to their physical size.

CTCCs have not been reported in blood for this MM.1S xenograft model previously. Moreover, we were able to identify only one previous direct report of clusters in the MM literature, where clusters were detected in patient blood samples (30). Therefore, we performed a secondary check for the presence of clusters in the blood by GFP fluorescence imaging of blood smears. These were taken from the second cohort of xenograft mice (C2) that were euthanized 36 days after injection. Example fluorescence micrograph images are shown in ***figs. 4g-r.*** Although the vast majority of observed MM.GFP+ cells on the bloods smears were individual CTCs (***fig. 4g-i***), we did observe a small number CTC clusters of varying size from a fewer than 10 (***fig 4j-o***), to dozens or hundreds of cells (***fig. 4p-r***). The blood smear imaging data did not permit accurate estimation of the numbers of CTCs in clusters, but the observed physical sizes were in good general agreement with our DiFC-derived estimates (***fig. 4f***). As such, we verified that MM.1S does in fact shed in clusters in this xenograft model, to our knowledge for the first time.

### 3.5 Estimation of CTC Burden from DiFC Measurements

Finally, we were interested in calibrating our DiFC measurements to the actual CTC burden. First, we calculated the estimated ‘corrected’ equivalent DiFC count rate per minute as shown in ***figure 5a***. To do this, we summed the contributions from individual cells (***fig. 3f***) with the contributions from the detected CTC clusters, which were derived from the cluster size estimates.

**Figure 5.**
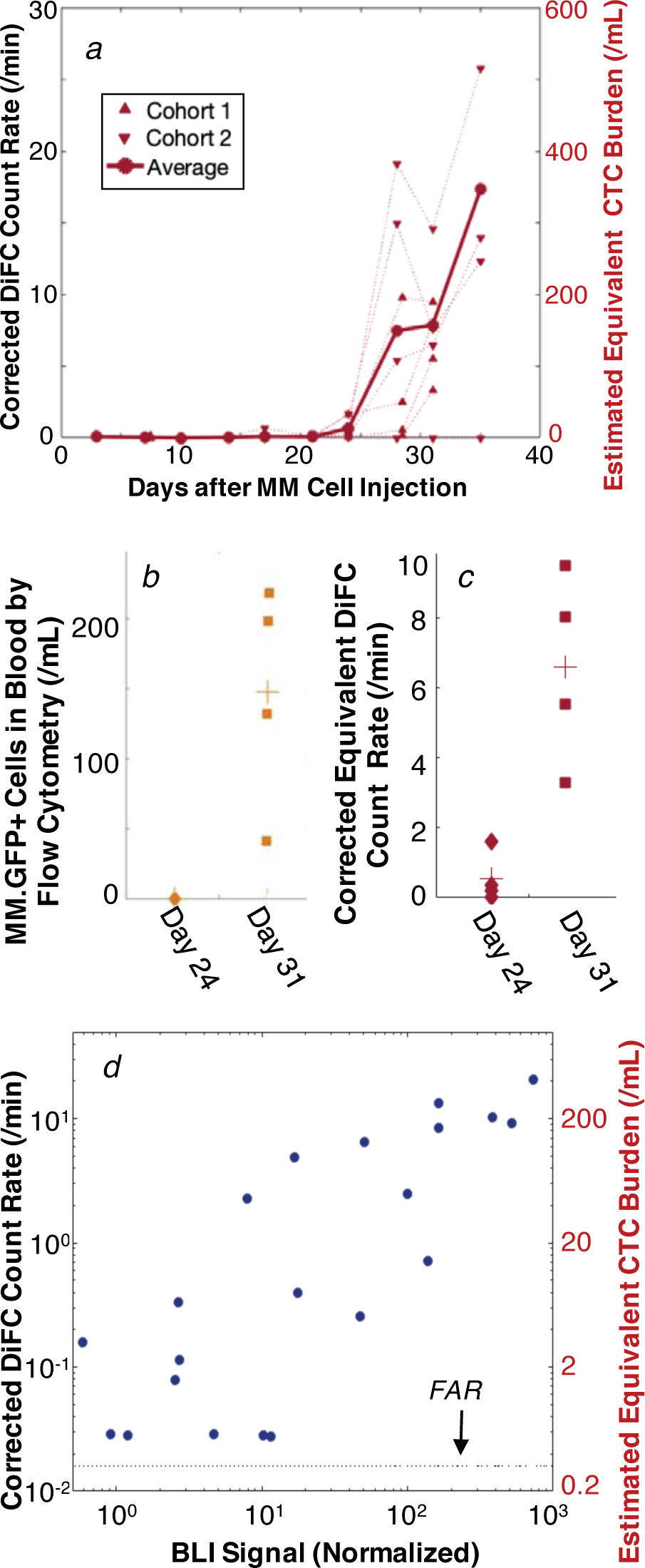
(a) we calculated the ’corrected’ DiFC count rate, which combined contributions from single CTCs and CTCCs (see text for details). (b) we drew blood samples on days 24 and 31 and counted MM GFP+ cells in the blood. (c) We compared these to the DiFC count rate on the same days, allowing us to estimate the detection sensitivity of DiFC. (d) the DiFC count rate also linearly correlated with MM disease burden measured by BLI for all mice and time-points (r^2^ = 0.75). All indicated points are above the false alarm rate of the system (black dotted line).

We then estimated the CTC burden in the peripheral blood (in cells per mL) from our measured DiFC count rate (in counts per minute) in two ways as follows. First, we used the average cell speed in the tail artery measured by DiFC of *v*_*s*_ = 26.5 mm/s. Making the simplifying assumption that the tail artery was a cylinder with 250 μm diameter with homogenous flow, this implies that the blood flow rate through DiFC in the artery was about 1.4 µL/s or 88 µL/min.

Second, in one cohort of mice (C1) we drew 200 µL of blood on day 24, and 0.5-1 mL of blood on day 31 (terminal). GFP-labeled MM cells in the blood samples were counted by flow cytometry (FC) as shown ***fig. 5b.*** On day 24, we were unable to detect any GFP labeled cells in the blood samples by FC whereas on day 31, small numbers of GFP+ CTCs were detectable. The average number of GFP+ cells in the 4 mice studies was 147 cells per mL of blood. The multiple steps of blood sample preparation including centrifugation and RBC lysing are expected to break up any CTC clusters in the blood. We also showed previously that this method of enumerating CTCs does not cause measurable loss of CTCs due to handling (18). We compared the estimates of CTC numbers in the blood to the corrected equivalent DiFC count rate from the same mice on the same days (***fig. 5c***), which was on average 6.6 counts/minute. Dividing the two yielded an estimated sampling rate of 45 µL/min.

The two methods of estimation were in disagreement by a factor of approximately two, which is reasonable considering the substantial uncertainty involved in both. It is possible that our method for estimating CTCs in blood from DiFC (including our threshold and signal processing method) under-estimates the true burden in blood. It is also possible that DiFC does not detect the most weakly-labeled GFP+ cells. Refining our signal processing algorithm, and better quantifying this effect is an ongoing area of work in our group. However, we chose a conservative (round) value of 50 µL/min for estimation of the absolute CTC burden on the right axis of ***fig. 5a***. This is lower than the previous sampling rate we estimated with our NIR DiFC system in nude mice (18), which we attribute to the smaller tail in SCID mice and correspondingly lower measured blood flow speed. Compared to other IVFC methods, this high blood sampling rate makes DiFC particularly well suited to studying rare circulating cells in the range of fewer than 100 cells per mL.

In addition, the overall MM tumor burden in the mice measured by BLI overall correlated well with the DiFC count rate as shown in ***fig. 5d*** (r^2^ = 0.75). In other words, while there was significant variability observed between mice in ***figures 2-4***, the same variability was measured on both systems for specific mice at specific times. This underscores the value of longitudinal study of individual mice, even in genetically identical mice with cultured cell lines.

## 4. Discussion and Conclusions

We previously reported DiFC (18) and showed that it was capable of detecting extremely rare circulating cells directly *in vivo* without having to draw blood samples. In this work, developed of a GFP-compatible DiFC system, and demonstrated its first use with an MM xenograft model in mice. As we noted previously, the detection sensitivity of blue-green DiFC – with respect to the minimum detectable fluorescent-labeling of target cells – is expected to be lower than microscopy-IVFC which uses confocal light detection (8). However, in ongoing studies in our lab, we have tested a number of GFP-labeled cell lines obtained from commercial and academic sources which thusfar are sufficiently bright for detection with DiFC.

To re-iterate, the main advantage of DiFC versus microscopy-IVFC methods is the use of diffuse light, which allows sampling of at least an order of magnitude greater circulating blood volumes. In this model, we estimated the sampling rate of 50 μL per minute, whereas microcopy IVFC samples on the order of 1 μL per minute. Prior work using microscopy-IVFC to study CTCs in an orthotopic tumor model in mice showed detection of a few cells per *hour* (31) or at concentrations in the range of 10^4^ cells per mL (32).

In contrast, in this study, the average count rate on the first day of detection (day 17 or 21, depending on the mouse) was 0.2 counts/min, which is equivalent to an estimated CTC burden of only 1 cell/mL of blood. The lowest count rates above the FAR (e.g. 0.03 counts/min for M3C1, day 24), was equivalent to approximately 0.4 cells/mL or about 1 cell in the entire peripheral blood volume of the mouse. These results demonstrate that DiFC is able to detect extremely rare circulating cells at the earliest stages of disease growth.

Such low CTC burdens are difficult to detect with liquid biopsy techniques involving non-terminal drawn blood samples. As illustrated in ***fig. 2i***, we also observed significant variability in the CTC count rate in short-term (minute-to-minute) intervals. Consideration of the count rate in a short 60s-time interval is analogous to estimating the CTC burden from a small (~50 μL) blood sample, which in this case could vary by more than a factor of 3 in the span of only 35 minutes. This was observed over all mice we studied with DiFC in these experiments. The short-term kinetics and statistics of CTC detection are the subject of ongoing work in our group.

The detection of CTC clusters in this MM xenograft model with DiFC was not anticipated based on our previous work with IVFC in this model, but was however not surprising, given that MM has been shown in a small number of reports to form clusters in the bone marrow (23,24) and in at least one previous report in patient blood (30). In addition, CTC clusters have been reported for many other cancer types. However, very little is known about MM CTCCs, such as their composition, frequency, size, and their short and long-term kinetics *in vivo*. As others have noted, this in part because most of methods of CTC isolation are not designed to capture clusters (15). While we interpret the data from the xenograft model with some caution, there were nevertheless some interesting features of our data:

- Clusters were consistently detected in circulation almost as soon as individual CTCs were observed in circulation (on day 21-24), suggesting that CTCCs shed with approximately the same kinetics.
- Clusters were observed with about 10% of the frequency of individual CTCs. This is in good general agreement with other literature reports of CTCCs, such as in breast cancer (17) and suggests that while CTCCs are rare, they are also not uncommon. The ratio of CTCs to CTCCs was approximately consistent throughout the studies.
- Most CTCCs were smaller than 10 cells (according to both DiFC and microscopy imaging estimates of size). A literature search failed to reveal other published estimates of clusters sizes specifically for MM.1S, or for MM in general. However these values are in good agreement with literature studies of CTCC cluster sizes for other cancers in mice and in humans (15).
- Clusters appeared to move very slowly relative to single CTCs in circulation, implying that they may experience greater adhesion forces along the blood vessel endothelium as others have suggested (15). Cluster signals on DiFC were consistently much wider than individual CTCs - the average pulse-width of single CTCs was ~39 ms, whereas clusters were on average 320 ms, and in individual cases longer than a second. The transit time between DiFC channels was several seconds or even minutes. This corresponds to a flow speed of a few mm per second or less, compared to tens of mm/s for single CTCs.

Overall, DiFC is a powerful small animal research tool that can provide useful complementary information about *in vivo* CTC and CTCC behavior for liquid biopsy assays. For example, kinetics of CTC and CTCC shedding can be measured with DiFC to inform the timing of blood draws, so that more detailed molecular and genetic analysis of cells can be performed. As we have noted previously, although DiFC can potentially be applied in larger limbs and species, it also relies on the use of fluorescence contrast which makes translation to human use challenging. Therefore, we view DiFC primarily as a small animal research tool in the foreseeable future.

Ongoing work in our group is exploring the use of the instrument in other models of animal models of metastasis. We are also studying the use of DiFC with targeted fluorescent molecular contrast agents, such as folate-targeted probes that have been shown to have high-affinity and specificity for CTCs (33). Moreover, we anticipate that DiFC could have significant utility with other studies involving rare circulating cell populations.

## Availability of Materials and Data

The datasets generated and analyzed during the current study, as well as the analysis code (written in Matlab) are available from the corresponding author on reasonable request.

## Acknowledgements

This work was funded by the National Institutes of Health (R01HL124315; NHLBI). We thank Eric Marple of EMVision LLC for helpful advice and discussion related to the fiber probe design. We thank Prof. Qianqian Fang (Northeastern University), for assistance using the MCX Monte Carlo software package.

---

**Figure S1.**
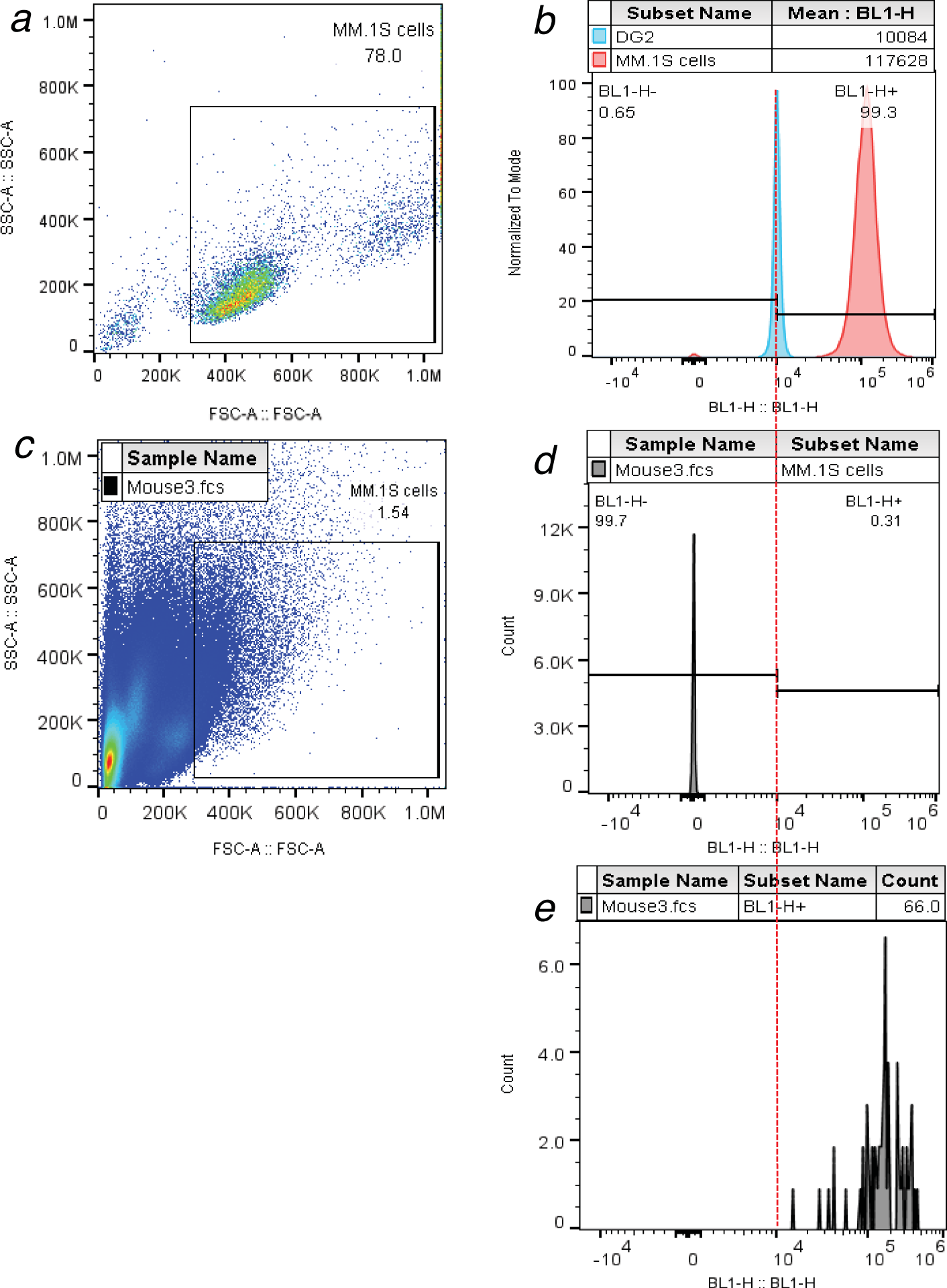
The FC gating strategy for counting MM.1S.GFP.Luc cells is shown. (a) SSC-FSC plot for MM.1S cells in culture. (b) the blue (BL1) fluorescence of MM.1S.GFP.Luc cells and dragon green 2 (DG2) microspheres. The mode intensity of DG2 microspheres was used as counting threshold since it was lower than cultured MM.1S.GFP.Luc cells. (c) SSC-FSC for a blood sample drawn from a mouse, with gate shown. RBCs were first depleted using a lysate protocol as described in the supplemental methods. (d) BL1 fluorescence signal in the blood. Most peaks are low-fluorescence debris or unlabeled cells. (e) Histogram of cells exceeding the DG2 threshold, yielding the cell counts in the sample.

